# A reduced vernalization requirement is a key component of the early-bolting trait in globe artichoke (*Cynara cardunculus* var. *scolymus*)

**DOI:** 10.1101/2024.01.23.576919

**Authors:** Rick Berentsen, Reyes Benlloch, Peter Visser, Francisco Madueño, Vicente Balanzà

## Abstract

Early-bolting is a major breeding objective for globe artichoke (*Cynara cardunculus* var. *scolymus* L.). It has been suggested that globe artichoke bolting time is linked to a vernalization requirement, although environmental conditions under which vernalized plants and controls have been grown may not always allow for proper comparison. Here, we defined morphological markers to monitor the vegetative-to-reproductive phase transition at the shoot apex and linked these to expression changes of homologues of key Arabidopsis flowering regulators *SOC1*, *FUL*, and *AP1*. Importantly, we developed an experimental setup where control and vernalized plants grow under comparable conditions. These tools together allowed for comparison of the vegetative-to-reproductive phase transition between early- and late-bolting genotypes and how they respond to vernalization. Our results show that vernalization requirement is significantly lower in early-bolting genotypes, supporting the view that the early-bolting trait is partly underlain by alterations in the network controlling vernalization response.

## Introduction

Globe artichoke [*Cynara cardunculus* L. var. *scolymus* (L.) Fiori] is thought to be a domesticated form of wild cardoon [*C. cardunculus* var. *sylvestris* (Lamk) Fiori] and cultivated for its sizable, closed, and compact inflorescences, which are colloquially referred to as “heads”. In Mediterranean climate zones cultivation cycles run from transplanting in summer till harvest between late autumn and late winter. Traditional varieties are often vegetatively propagated whereas modern hybrids are propagated by seeds and designed to be sown and harvested within a single cultivation cycle. Although seeded hybrids have demonstrated their potential for superior yield, quality, and uniformity over traditional varieties, they tend to produce later than many early producing traditional varieties ^[1][2][3]^ ^[4]^. For that reason, combining the superior quality and production of hybrids with the earliness of traditional varieties is a major objective for globe artichoke breeders.

Globe artichoke has a seasonal life cycle. At the start of the season, one or more rosettes develop from the roots. These embody the vegetative phase which is characterized by the continuous production of leaves by the shoot apical meristem (SAM), without elongation of the stem internodes. During the vegetative-to-reproductive phase transition, which is also known as the floral transition, the hitherto leaf-producing SAM is converted into an inflorescence meristem (IM), that gives rise to inflorescence stems and capitulae (heads). Developmental stages in cardoon and globe artichoke have been described and classified ^[5][6][7][8]^. These scales however do not describe in detail the changes that occur in the shoot or inflorescence apex between the vegetative-to-reproductive phase change and bolting. Moreover, molecular markers for the vegetative-to-reproductive phase transition, potentially useful for studies on the genetic architecture of flowering, are not available for globe artichoke.

To enable successful reproduction in plants, the vegetative-to-reproductive phase needs to be precisely timed in accordance with a variety of endogenous and environmental cues such as age, photoperiod, vernalization, ambient temperature or drought ^[9][10][11][12]^. These flowering cues are sensed by flowering inductive pathways that converge upon a set of flowering integrators that finally direct the vegetative-to-reproductive phase transition. These pathways form an intricate genetic network, which has been extensively studied and reviewed in the model species Arabidopsis (*Arabidopsis thaliana*) and cereals such as rice, barley, and wheat ^[9][13]^ ^[14][15]^, although the relevance of the different pathways and mechanisms controlling flowering in non-model species are often poorly understood.

In Arabidopsis the major flower inductive pathways are the vernalization, photoperiod, age, gibberellin, and autonomous ones ^[16]^. These pathways converge on specific floral integrators. The photoperiod pathway enhances the expression of the floral integrator *FLOWERING LOCUS T* (*FT*), a phosphatidylethanolamine binding protein (PEBP) family member ^[17][18]^ that positively controls the expression of MADS box floral integrator gene *SUPPRESSOR OF OVEREXPRESSION OF Constans 1* (*SOC1*) ^[19]^ ^[20]^ and *FRUITFULL* (*FUL*) ^[21][22]^. *SOC1* also functions independently from *FT*, being activated directly by the vernalization and autonomous pathways through the repression of the MADS box gene *FLOWERING LOCUS C* (*FLC*) ^[23]^. Moreover, *SOC1* and *FUL* are activated by the age pathway through the microRNA miR156-mediated activity of different *SQUAMOSA PROMOTER BINDING PROTEIN-LIKE* genes (*SPL3, SPL9*, and others) ^[24]^ and by the gibberellin pathway that promotes *SOC1* and *FUL* expression through the degradation of DELLA proteins and the release of active SPL proteins, among others ^[25][26][27]^. Finally, *SOC1* and *FUL* function in a partially redundant manner to control the expression of the meristem identity gene *LFY* ^[28][29][30]^.

Different studies have indicated that individual genes, or gene families, that constitute this network are largely conserved between Arabidopsis and Asteraceae like lettuce (*Lactuca sativa* L.), chicory (*Cichorium intybus* L.), safflower (*Carthamus tinctorius* L.), Chrysanthemum (*Chrysanthemum* sp.) and gerbera (*Gerbera hybrida* L.), although gene orthology and function cannot always be inferred directly ^[31][32][33][34][35][36][37]^. Due to their central role in floral signal integration and floral meristem and organ identity, members of the MADS box gene family have been studied in Asteraceae and homologs of *SOC1*, *FUL* and *AP1* have been reported for chrysanthemum, gerbera, lettuce, safflower and sunflower ^[31][35][37][38][39]^ ^[40][41]^. Despite this, knowledge about the identity of the floral integrators in globe artichoke is scarce, as well as about cues that trigger the floral transition process.

With respect to photoperiod, different authors have considered globe artichoke to be either an obligate long-day plant ^[42]^, a short-day plant ^[43]^, or considered it to flower independent of photoperiod ^[44]^, suggesting a genotype dependent response. More consistent links have been found with gibberellic acid (GA) and vernalization. Application of GA3 is known to advance the moment of bolting in globe artichoke ^[45][46][47]^. Vernalization, a predetermined requirement for cold exposure to acquire flowering competence, has been reported to be a major determinant of flowering in globe artichoke ^[44]^ ^[48]^. Moreover, it has been suggested that the presence of a vernalization requirement is linked to late-bolting genotypes, bolting after the winter, whereas the absence of such a requirement might explain the existence of genotypes that can bolt before winter ^[42][43]^. However, most studies on the effect of, or requirement for, vernalization in globe artichoke have been performed either under non-controlled field conditions ^[48]^ ^[43]^, by exposing plants to vernalizing temperatures only during part of their life cycle ^[49]^ ^[50][51]^, or do not consider the time of bolting in individual plants ^[48]^. This means that, although these studies provide valuable information on the role of vernalization regimes for improved cultivation practices, their results do not allow to quantify the precise effect of vernalization on the time of bolting and/or to compare results between studies.

In order to better understand the floral transition in globe artichoke we have characterized the morphological changes that occur at the macroscopic and microscopic level in the shoot apex that are associated with the vegetative-to-reproductive phase transition. We also established an experimental procedure that enables study of the vernalization requirement of globe artichoke under controlled and comparable conditions, as well as identified homologs of key regulators of the floral transition in Arabidopsis such as *SOC1*, *FUL* and *AP1* for globe artichoke that can be used as molecular markers for floral transition. These tools have allowed us to compare the vernalization requirement between early- and late-bolting genotypes, indicating that all genotypes under study are able to respond to vernalization. Late-bolting genotypes however respond comparatively stronger to vernalization and have a higher requirement for vernalization than early-bolting ones.

## Results

### Novel morphological markers of the shoot apex associated with the transition to the reproductive phase

The different developmental stages during the vegetative-to-reproductive phase transition in artichoke have been described at the macroscopic level. Bolting stage 0 has been proposed to define plants with no signs of bolting, whereas bolting stage A is assigned to the time when the primary inflorescence is palpable in the center of the basal leaf rosette ^[5]^ ^[8]^. In order to determine the moment of initiation of vegetative-to-reproductive phase transition in artichoke we decided to complement the previously proposed developmental stages ^[5]^ ^[8]^ at the microscopic level. For this, we dissected inflorescence apices and observed them both macro- and microscopically in weekly intervals. We defined five new stages that describe the morphological changes in the apex during the time before the primary inflorescence becomes palpable inside the rosette (Supplementary Table S1). We named these five new stages “pre-bolting stage 0” to “pre-bolting stage 4”. The first detectable sign of the vegetative-to-reproductive phase transition having been initiated is a change in the shape of the hitherto globose or flat vegetative apex, i.e., pre-bolting stage 0 (Figure 1 A-B), towards a domed, and later pointed or triangular form, which we designated “pre-bolting stage 1” (Figure 1 C-D). Pre-bolting stage 2 starts with the elongation of the inflorescence meristem and the formation of the first bracts of the primary inflorescence at the tip of the inflorescence apex (Figure 1 E-F). Subsequently, pre-bolting stage 3 is characterized by an increased number of bracts and further elongation of the primary inflorescence, which now reaches about 1 cm in size (Figure 1 G). Pre-bolting stage 4 (Figure 1 H) commences with the initiation of second order inflorescences and coincides with bolting stage A, during which the primary inflorescence becomes palpable in the center of the rosette (Figure 1 I) ^[8]^.

**Figure 1:**
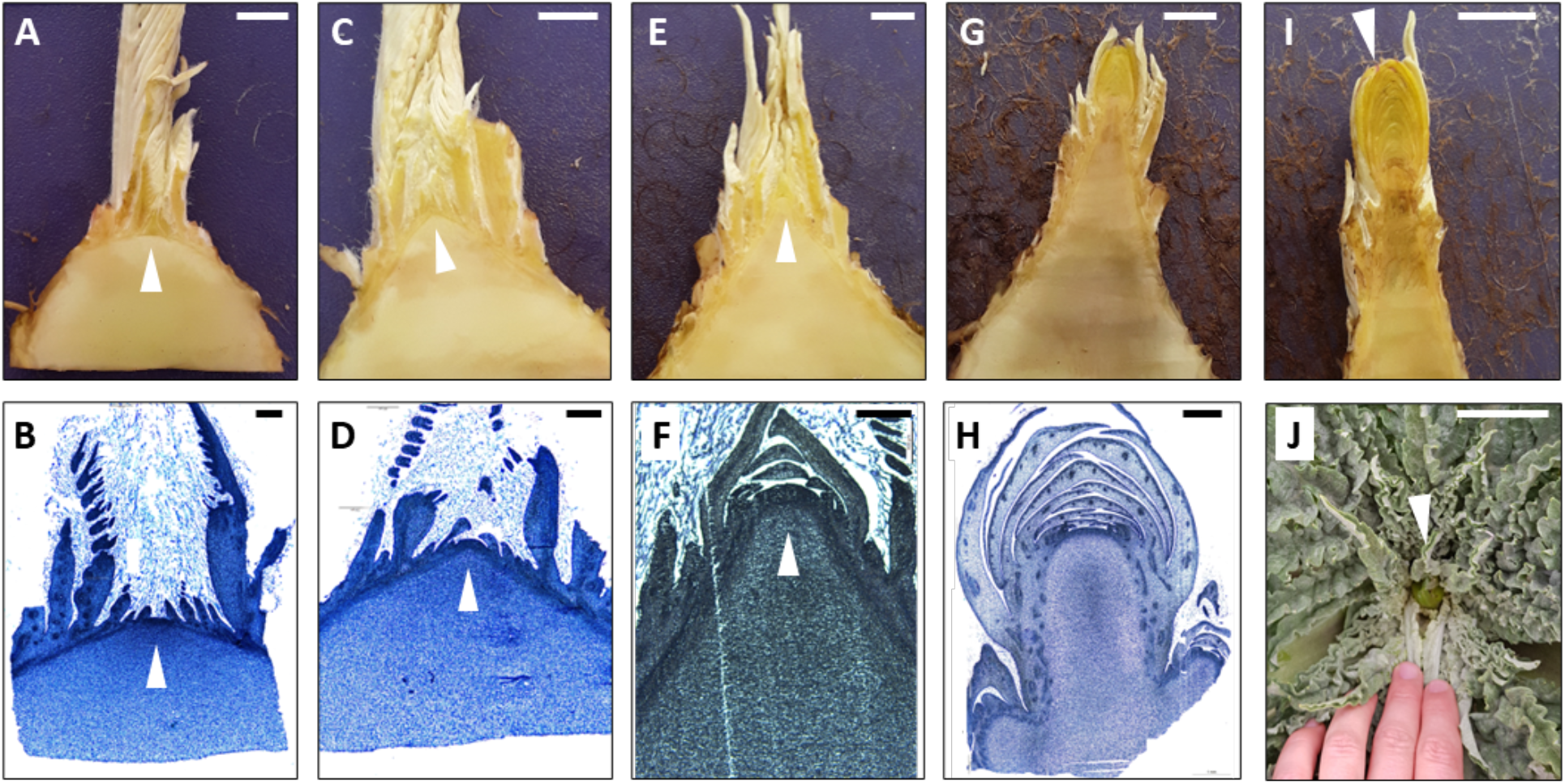
Morphological changes in the SAM around the moment of vegetative-to-reproductive phase transition and a classification of developmental stages. **A.** pre-bolting stage 0 (genotype c20, 87 days post transplanting (d.p.t.)), **B.** pre-bolting stage 0 (genotype CARI, 127 d.p.t.), **C.** pre-bolting stage 1 (c20, 123 d.p.t.), **D.** pre-bolting stage 1 (CARI, 126 d.p.t), **E**. pre-bolting stage 2 (c20, 130 d.p.t.), **F**. pre- bolting stage 2 (‘Green Queen F1’, 140 d.p.t.), **G.** pre-bolting stage 3 (c20, 144 d.p.t.), **H.** pre-bolting stage 4 (CARI, 147 d.p.t.), I. bolting stage A (CARI, 144 d.p.t.), J. bolting stage “B” for comparison. Triangles point to SAM (A-D), the primary inflorescence primordium (E-F), or emerging primary capitulum (I-J). Scale bars: 500 μm (A-G), 1000 μm (H-I), 4 cm (J).

### Early-bolting is linked to a reduced vernalization requirement

Genetic control of bolting time in globe artichoke is poorly known. Some studies have proposed a relevant role for vernalization as a determinant of bolting time in different genotypes ^[42]^ ^[48]^. However, conclusions from these studies require assessment because of non-controlled conditions, i.e. comparing genotypes grown in different time frames, climate zones, or at different latitudes. Thus, most experiments have been performed in the field under conditions in which the individual contributions of vernalization and photoperiod cannot easily be assayed as separate variables ^[43]^. In other studies, an artificial vernalization treatment was applied exclusively during an early developmental phase, after which plants were grown and observed under field conditions ^[49]^ ^[50]^ ^[51]^. In such cases it is not possible to rule out the possible contribution of other factors, such as devernalization, referring to a possible reversal of the vernalized state after prolonged exposure to temperatures above a certain threshold ^[52]^.

To overcome these limitations and determine the effect of vernalization on the time of bolting in early- and late-bolting genotypes under appropriate conditions, we designed an experimental setup where half of the plants were grown in a net house in a standard growing cycle (transplanting near the end of September and anthesis by the end of April), which naturally includes a vernalization period. The remainder plants were grown under the same conditions, in a net house adjacent to the aforementioned one, but where vernalizing temperatures (<12°C) were prevented by means of a thermostat-controlled heating system (Supplementary Figure S1). The genotypes used in the experiments were two early-bolting clones (“c1” and “c70”), two late-bolting clones (“c20” and “c154”), two inbred early-bolting lines (“VESB” and “VER”) and one inbred late-bolting line (“CARI”).

The experiments were carried out during three seasons, between 2019 and 2022, comparing plants grown with or without vernalization. The genotypes studied were characterized during at least two seasons, except the inbred line “VER” which was only characterized during the first season. For the vernalized plants, the number of chilling hours (i.e., hours <10°C) was 698, 837, and 832, in the three seasons respectively (Supplementary Table S2). Plants were observed weekly, and their developmental stage scored by eye. A plant was considered to have bolted upon reaching bolting stage B (Figure 1J and Supplementary Table S1).

The results of the experiments in the three seasons are summarized in Table 1. Out of the 1,170 plants analyzed, a total of 55 did not bolt at the end of the season, 44 that were non-vernalized and 11 that were vernalized. Out of the 44 non-vernalized plants that remained in the vegetative phase, 6 were from early-bolting genotypes and the remainder 38 from late-bolting genotypes, more specifically c20 (7), c154 (7) and CARI (24). This suggests that, in the absence of vernalizing temperatures, late-bolting genotypes have a higher propensity towards remaining vegetative at the end of the season than the early-bolting ones. Under vernalization conditions, at the moment of bolting, early-bolting genotypes c1, c70, VESB, and VER, had accumulated an average of 604.0, 403.8, 493.1 and 476.8 hours below 10°C respectively, whilst the late-bolting genotypes (c20, c154, and CARI) accumulated considerably more hours below 10°C (777.5, 773.3, and 737.0 respectively) (Table 1). Interestingly, late-bolting genotypes accumulated similar amounts of hours below10°C.

**Table 1.** Proportion of plants bolted in time per genotype, season, and treatment.

| earliness | genotype | treatment | season | n_obs<sup>a</sup> | not bolted<sup>b</sup> | mean date bolting stage B<sup>c</sup> | DTB |  | mean daylength<sup>d</sup> | accumulated hours <10°C |  |
| --- | --- | --- | --- | --- | --- | --- | --- | --- | --- | --- | --- |
|  |  |  |  |  |  |  | mean | sd |  | per season | mean<sup>e</sup> |
| early | c1 | no vernalization | 2019-2020 | 19 | 0 | 2020-02-15 | 144.9 | 7.8 | 10.8 | 1.5 | 0.9 |
| early | c1 | no vernalization | 2020-2021 | 15 | 1 | 2021-02-08 | 137.1 | 3.4 | 10.5 | 0.7 |  |
| early | c1 | no vernalization | 2021-2022 | 57 | 2 | 2022-02-14 | 137.2 | 6.5 | 10.8 | 0.5 |  |
| early | c1 | vernalization | 2019-2020 | 0 | 0 | NA | NA | NA | NA | NA | 604.0 |
| early | c1 | vernalization | 2020-2021 | 18 | 0 | 2021-02-09 | 138.8 | 3.0 | 10.6 | 617.8 |  |
| early | c1 | vernalization | 2021-2022 | 59 | 0 | 2022-02-06 | 129.9 | 4.6 | 10.5 | 590.3 |  |
| early | c70 | no vernalization | 2019-2020 | 32 | 0 | 2020-01-09 | 107.8 | 8.5 | 9.8 | 1.5 | 0.7 |
| early | c70 | no vernalization | 2020-2021 | 18 | 0 | 2021-01-19 | 117.1 | 10.8 | 10.0 | 0.1 |  |
| early | c70 | no vernalization | 2021-2022 | 57 | 3 | 2022-01-16 | 108.1 | 10.4 | 9.9 | 0.5 |  |
| early | c70 | vernalization | 2019-2020 | 0 | 0 | NA | NA | NA | NA | NA | 403.8 |
| early | c70 | vernalization | 2020-2021 | 13 | 1 | 2021-01-23 | 121.4 | 10.8 | 10.1 | 540.8 |  |
| early | c70 | vernalization | 2021-2022 | 52 | 0 | 2022-01-05 | 97.1 | 10.0 | 9.7 | 266.7 |  |
| early | VESB | no vernalization | 2019-2020 | 57 | 0 | 2020-02-05 | 134.5 | 12.3 | 10.5 | 1.5 | 1.0 |
| early | VESB | no vernalization | 2020-2021 | 0 | 0 | NA | NA | NA | NA | NA |  |
| early | VESB | no vernalization | 2021-2022 | 65 | 0 | 2022-02-17 | 140.4 | 7.4 | 10.9 | 0.5 |  |
| early | VESB | vernalization | 2019-2020 | 51 | 0 | 2020-01-21 | 119.4 | 12.6 | 10.0 | 431.6 | 493.1 |
| early | VESB | vernalization | 2020-2021 | 0 | 0 | NA | NA | NA | NA | NA |  |
| early | VESB | vernalization | 2021-2022 | 49 | 0 | 2022-02-02 | 125.4 | 5.8 | 10.4 | 554.7 |  |
| early | VER | no vernalization | 2019-2020 | 53 | 0 | 2020-02-12 | 141.4 | 7.7 | 10.7 | 1.5 | 1.5 |
| early | VER | no vernalization | 2020-2021 | 0 | 0 | NA | NA | NA | NA | NA |  |
| early | VER | no vernalization | 2021-2022 | 0 | 0 | NA | NA | NA | NA | NA |  |
| early | VER | vernalization | 2019-2020 | 51 | 0 | 2020-01-26 | 124.5 | 14.3 | 10.2 | 476.8 | 476.8 |
| early | VER | vernalization | 2020-2021 | 0 | 0 | NA | NA | NA | NA | NA |  |
| early | VER | vernalization | 2021-2022 | 0 | 0 | NA | NA | NA | NA | NA |  |
| late | c20 | no vernalization | 2019-2020 | 0 | 0 | NaN | NA | NA | NA | NA | 48.2 |
| late | c20 | no vernalization | 2020-2021 | 8 | 0 | 2021-03-19 | 176.6 | 9.1 | 12.1 | 28.0 |  |
| late | c20 | no vernalization | 2021-2022 | 5 | 7 | 2022-04-12 | 199.6 | 3.1 | 13.0 | 68.5 |  |
| late | c20 | vernalization | 2019-2020 | 0 | 0 | NA | NA | NA | NA | NA | 777.5 |
| late | c20 | vernalization | 2020-2021 | 10 | 4 | 2021-03-15 | 172.0 | 0.0 | 11.9 | 723.5 |  |
| late | c20 | vernalization | 2021-2022 | 12 | 0 | 2022-03-15 | 168.9 | 3.6 | 11.9 | 831.5 |  |
| late | c154 | no vernalization | 2019-2020 | 0 | 1 | NA | NA | NA | NA | NA | 57.4 |
| late | c154 | no vernalization | 2020-2021 | 10 | 1 | 2021-03-29 | 191.8 | 12.6 | 12.5 | 51.9 |  |
| late | c154 | no vernalization | 2021-2022 | 25 | 5 | 2022-04-10 | 194.3 | 5.1 | 12.9 | 62.9 |  |
| late | c154 | vernalization | 2019-2020 | 0 | 0 | NA | NA | NA | NA | NA | 773.3 |
| late | c154 | vernalization | 2020-2021 | 10 | 0 | 2021-03-12 | 169.9 | 3.4 | 11.8 | 715.0 |  |
| late | c154 | vernalization | 2021-2022 | 26 | 4 | 2022-03-15 | 171.4 | 3.0 | 11.9 | 831.5 |  |
| late | CARI | no vernalization | 2019-2020 | 16 | 15 | 2020-03-29 | 211.4 | 20.6 | 12.5 | 1.5 | 20.8 |
| late | CARI | no vernalization | 2020-2021 | 0 | 0 | NA | NA | NA | NA | NA |  |
| late | CARI | no vernalization | 2021-2022 | 39 | 9 | 2022-04-04 | 189.7 | 10.0 | 12.7 | 40.0 |  |
| late | CARI | vernalization | 2019-2020 | 32 | 0 | 2020-03-03 | 161.1 | 3.6 | 11.5 | 642.4 | 737.0 |
| late | CARI | vernalization | 2020-2021 | 0 | 0 | NA | NA | NA | NA | NA |  |
| late | CARI | vernalization | 2021-2022 | 43 | 2 | 2022-03-15 | 171.9 | 5.9 | 11.9 | 831.5 |  |

<sup>a</sup> nr of observations
<sup>b</sup> number of individuals that had not bolted at the end of the experiment
<sup>c</sup> mean date of reaching bolting stage B
<sup>d</sup> daylength refers to the mean calculated number of hours that the sun is above the horizon on a given day upon reaching bolting
stage B
<sup>e</sup> mean of cumulative hours over three seasons

We also observed that the plants from the early-bolting genotypes, both vernalized and non-vernalized, bolted between January and February, a time frame associated with short-day conditions (Table 1). In contrast, the late-bolting genotypes, under no-vernalization conditions, bolted around the end of March and early April, whereas under vernalization conditions bolting occurred around the middle of March. This time frame indicates that bolting in these genotypes is not strictly confined to short-day conditions (Table 1).

To measure the level of earliness we considered the number of days to bolting stage B (DTB). Then DTB were modelled using a linear mixed model to account for both fixed effects and for random effects such as between-plot variation between different seasons (random effects). The analysis revealed that the terms “genotype” (p-value = 1.16*10^-80^), “treatment“ (p-value = 3.01*10^-30^), and “genotype:treatment” (p-value = 9.09*10^-6^) significantly affected bolting time, whereas “season” did not (p-value = 0.23) (Supplementary Table S3). The estimated means were extracted from the model for each “genotype:treatment” combination, and the significance of differences between best linear unbiased estimates (BLUEs) was assessed by whether the difference between BLUEs was larger or smaller than the value of the least significant difference (LSD, 6.86 days) (Figure 2A). The statistical model showed that vernalization had a significant effect on bolting time in all genotypes that were observed in multiple seasons, indicating that both early- and late-bolting genotypes were sensitive to vernalization (Figure 2A).

**Figure 2:**
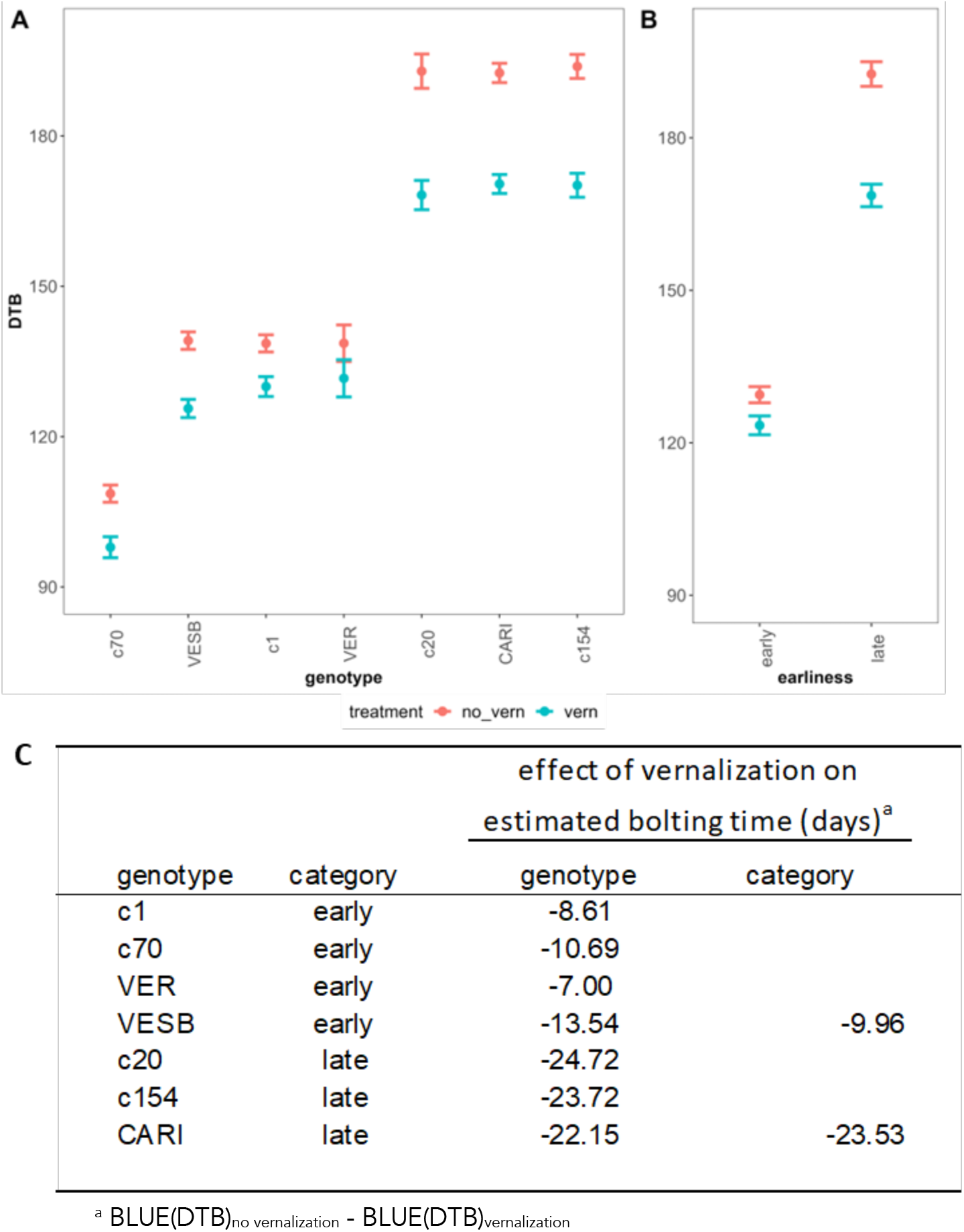
Model predictions and effects of vernalization on days to bolting (DTB) in early- and late-bolting genotypes. **A.** BLUEs (best linear unbiased estimates) for each genotype:treatment combination. Non bolters were not included in the analysis. Error bars = predicted interval. **B.** BLUEs categorized by early- vs late-bolting habit. **C.** Effect of vernalization on bolting time BLUEs per genotype and genotype category. LSD (least significant difference) for both models is 6.86.

To determine whether the effect of vernalization was different between the early- and the late-bolting genotypes, the modelling was repeated substituting explanatory variable “genotype” with “earliness”, a dichotomous variable distinguishing early-bolting genotypes (c1, c70, VESB and VER) from late-bolting ones (c20, c154 and CARI). In this model, the term “earliness” was significant whilst “season” was not (Supplementary Table S3). Differences in estimated means were not significant for the effect of vernalization on early-bolting genotypes (6.02 < 6.86) but they were significant for late-bolting genotypes (23.89 > 6.86) (Figures 2B and 2C). These results indicate that the late-bolting genotypes significantly respond to vernalization whereas the early-bolting genotypes, as a category, probably due to different vernalization responses inside the group, apparently do not respond.

Whereas in the late-bolting genotypes vernalization advanced the estimated mean bolting time by 23.53 days in the model, in the early-bolting ones vernalization advanced mean bolting time by 9.96 days (Figure 2C). More specifically, in the early-bolting genotypes the effect of vernalization was largest for VESB (13.54 days advance), followed by c70 (10.69 days), c1 (8.61 days) and VER (7.00 days), whilst in the late-bolting genotypes the effect was largest for c20 (24.72 days), followed by c154 (23.72 days) and CARI (22.15 days) (Figure 2C). The larger effect of vernalization on late-bolting genotypes is in agreement with the latter having accumulated more hours below 10°C, as registered during the vernalization experiment (Supplementary Table S2, Supplementary Figure 2), which indicates that early-bolting genotypes have a lower vernalization requirement for bolting than late-bolting genotypes. This analysis clearly indicates that our experimental setup allows for measuring the effect of vernalization in the tested genotypes under comparable conditions, making it feasible to extend the results of this study to the behavior of plants under field conditions.

Bolting is a late and advanced phase of the floral transition, which occurs days before the vegetative-to-reproductive phase change is detected. To better understand the effect of vernalization on the timing of the vegetative-to-reproductive phase transition in these plants we decided to monitor the transition in selected early and late-bolting genotypes by direct observation of the plant apices. For this purpose, we selected early-bolting genotype c1 and late-bolting genotype c154. During the 2020-2021 season we performed an additional vernalization experiment wherein shoot apices were collected and analyzed at different time points before bolting. The vegetative-to-reproductive phase transition was considered to have commenced when the shoot apical meristem reached pre-bolting stage 1 (Figure 1 A-B, Supplementary Table S1). This experiment showed that vernalized plants from genotype c1 reached this stage on average at 88.3 days after transplanting whereas non-vernalized plants reached this stage at 84.1 days after transplanting, having accumulated only 148.7 hours below 10°C (Table 2, Figure 3). This analysis indicates that vernalization apparently does not affect the time of floral transition of early-bolting genotype c1. In contrast, non-vernalized plants from late-bolting genotype c154 reached pre-bolting stage 1 later than plants from genotype c1, at an average of 131.6 days after transplanting (Table 2, Figure 3). Under vernalization conditions, plants from genotype c154 showed a significant advancement of the vegetative-to-reproductive phase transition with respect to non-vernalized plants (115.0 days, p=0.043), having accumulated 459.4 hours below 10°C (Table 2, Figure 3). This analysis also reflected that the two different genotypes characterized initiate the floral transition in short-day conditions (third week of December for both vernalized and non-vernalized c1plants, last days of January for non-vernalized c154 plants, and middle January for vernalized ones) (Table 2).

**Figure 3:**
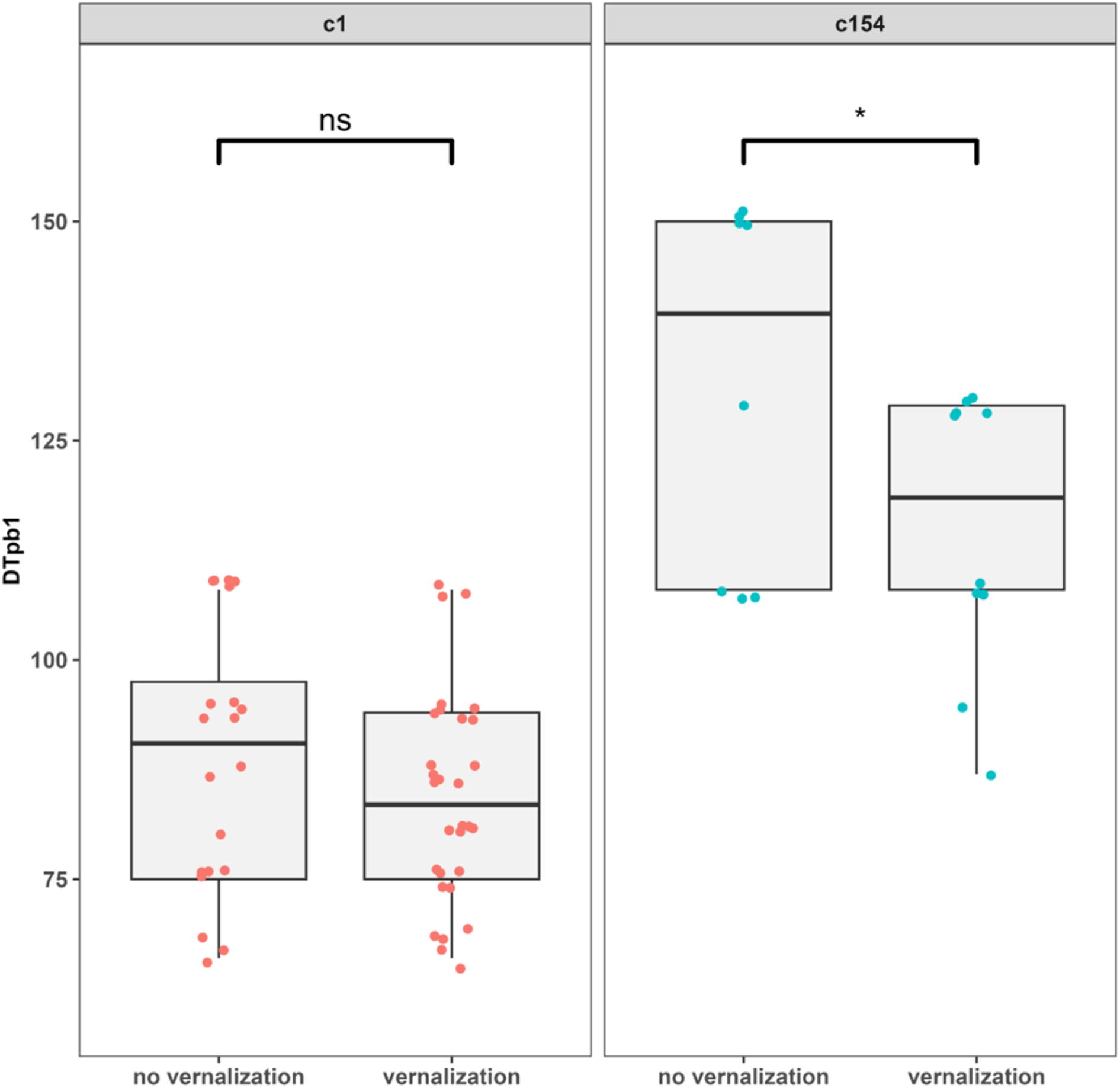
Days to pre-bolting stage 1 (DTpb1) in early-bolting genotype c1 and late-bolting genotype c154. Data from 2020-2021 season. Red and blue dots represent individual samples. Significance levels: “n.s.”, not significant, “*”, 0.01 < P < 0.05.

**Table 2.** Proportion of plants bolted in time per genotype, season, and treatment.

| earliness | genotype | treatment | season | n_obs<sup>a</sup> | mean date<br>bolting stage B<sup>c</sup> | DTpb1 |  | mean<br>daylength<sup>d</sup> | acc. hours<br><10°C<sup>e</sup> |
| --- | --- | --- | --- | --- | --- | --- | --- | --- | --- |
|  |  |  |  |  |  | mean | sd |  |  |
| early | c1 | no vernalization | 2020-2021 | 20 | 2020-12-21 | 88.3 | 15.0 | 9.7 | 0.0 |
| early | c1 | vernalization | 2020-2021 | 30 | 2020-12-17 | 84.1 | 12.0 | 9.6 | 148.7 |
| late | c154 | no vernalization | 2020-2021 | 9 | 2021-01-29 | 127.4 | 23.2 | 10.4 | 0.3 |
| late | c154 | vernalization | 2020-2021 | 10 | 2021-01-17 | 115.0 | 16.2 | 10.0 | 459.4 |

<sup>a</sup> nr of observations
<sup>b</sup> number of individuals that had not bolted at the end of the experiment
<sup>c</sup> mean date of reaching pre-bolting stage 1
<sup>d</sup> daylength refers to the calculated number of hours that the sun is above the horizon on a given day
<sup>e</sup> number of accumulated hours below 10°C

### Analysis of artichoke MADS-box genes to develop expression markers for the vegetative-to-reproductive phase transition

We have identified morphological markers in the shoot apex to assess the developmental stages during the transition from the vegetative to the reproductive phases in globe artichoke (Figure 1). In order to generate gene-expression markers for the vegetative-to-reproductive phase transition, which might signal developmental changes not revealed by morphological markers, we identified homologs of genes with known roles as flowering pathway integrators ^[53]^. MADS-box genes *SOC1* and *FUL* have been described as flowering integrators in multiple plant species ^[54]^ ^[34]^. In Arabidopsis, increases in expression levels of these genes accompany the vegetative-to-reproductive phase transition, during which they are strongly upregulated ^[14]^. Thus, we looked for homologues of the *SOC1* and *FUL* genes in the globe artichoke genome ^[55]^.

A BLASTp search of MADS box proteins from Arabidopsis, tomato and lettuce on the globe artichoke genome yielded a total of 82 hits. In plants, two major lineages of MADS box genes can be distinguished by phylogenetic studies, being Type I and Type II (MIKC-like). Type I genes contain a MADS domain, whereas Type II genes possess an additional Keratin-like domain (K-domain) ^[56]^. A Pfam domain scan of the 82 proteins identified significant SRF/MADS domains (PF00319.21) in 67 proteins and significant K- domains (PF01486.20) in 33 proteins. The genes were named *CcMADS1* till *CcMADS81* (Supplementary Table S4). After clustering the proteins with Arabidopsis, tomato, and lettuce homologs, they were divided into 28 type-I and 53 type-II MADS box genes (Supplementary Figure S3). In a total of 53 type II MADS box genes, we identified 3 MIKC*-type (M**δ** subfamily) genes. The remainder 50 MIKC^C^-genes were divided into 13 subfamilies. In brief, we identified 81 MADS box genes in the globe artichoke genome that represent the main clades found in other angiosperms. In a more detailed phylogenetic study of the SOC1 subfamily, we clustered globe artichoke proteins with SOC1 homologs and subfamily members from Arabidopsis, tomato, lettuce, safflower and gerbera (Figure 4A). Of the five globe artichoke *SOC1* subclade members, *CcMADS66*, *CcMADS68*, and *CcMADS72*, contain both an SRF and a K-box Pfam domain (Supplementary Table S4). Moreover, when considering both gene structure and protein sequence similarity to *AtSOC1, CcMADS68* and *CcMADS72* were most similar (50.9% and 63.9% respectively). These two genes also contain the canonical C-terminal “eVETeLvIGpP” *TM3*/*SOC1* motif ^[57]^, which was not found in the remainder genes except for *CcMADS66*. Taken together, these data suggest that the SOC1 subfamily contains at least two true *SOC1* homologues, *CcMADS72* and *CcMADS66,* which we designated *CcSOC1a* and *CcSOC1b*. Remainder homologs, *CcMADS78*, *CcMADS68* and *CcMADS15*, were designated *CcSOC1Like-A, CcSOC1Like-B*, and *CcSOC1Like-C*.

**Figure 4:**
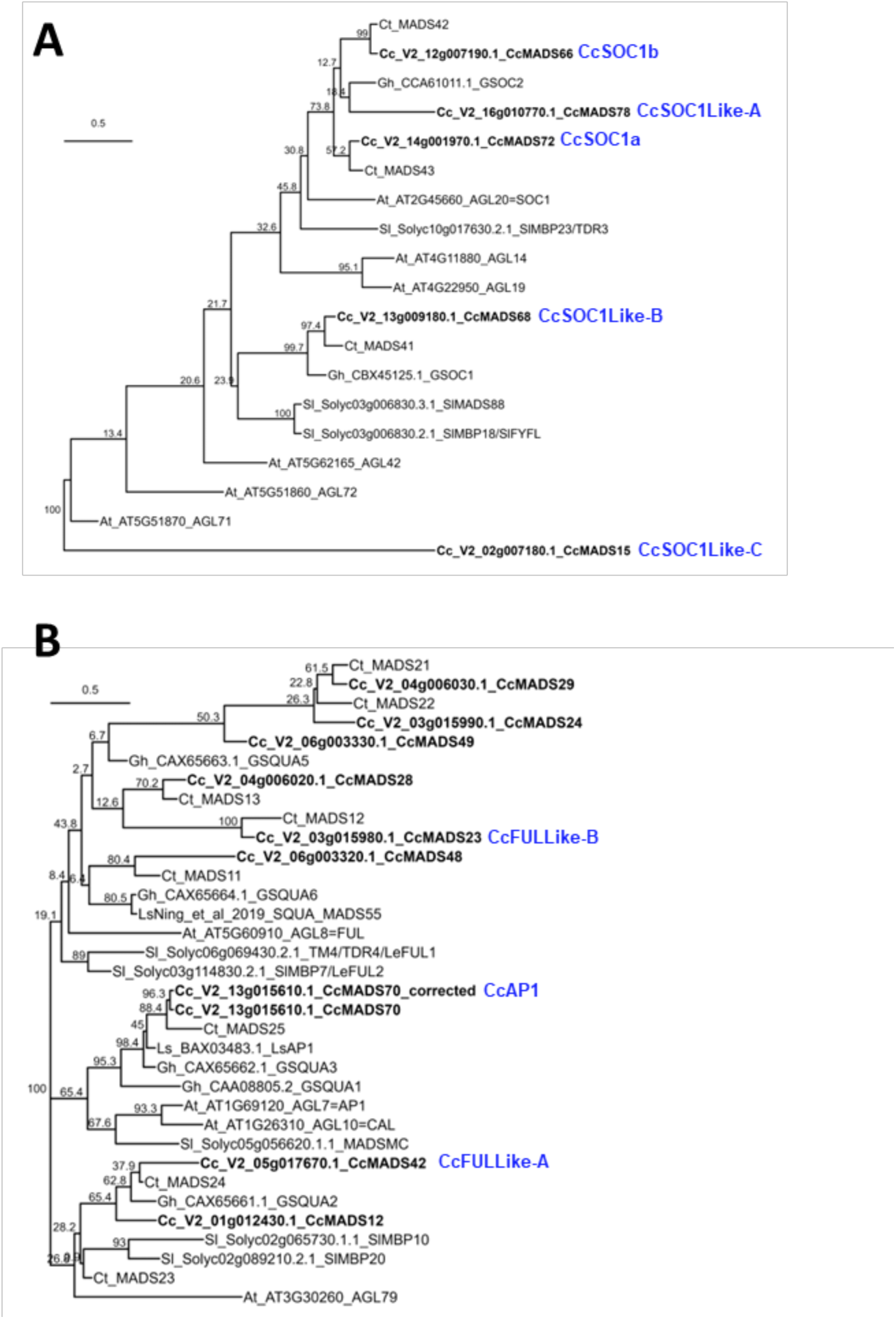
ML phylogenetic tree of SOC1 and AP1 proteins including globe artichoke. **A.** SOC1 proteins. **B.** AP1 proteins. Species are Cc = globe artichoke, Sl = tomato, At = Arabidopsis, Ct = safflower, Gh = gerbera. Nodes annotated with 1000-bootstrap values. Proteins from globe artichoke in bold italics.

To study the AP1/FUL subfamily in globe artichoke in more detail, proteins belonging to this subfamily, CcMADS23, CcMADS24, CcMADS28, CcMADS29, CcMADS42, CcMADS48, CcMADS49, and CcMADS70, were clustered with AP1 homologues from Arabidopsis, tomato, lettuce and safflower (Figure 4B). The globe artichoke proteins, ranging from 74 a.a. to 205 a.a. (Supplementary Figure S4) are all shorter than *AP1* from Arabidopsis (256 a.a.) or *LsAP1* from lettuce (253 a.a.) ^[41]^, albeit that short proteins in the *AP1* family are encountered in the lettuce as well ^[35]^. The number of exons in the globe artichoke *AP1* genes ranged from one exon (*CcMADS24*, *CcMADS29* and *CcMADS49*) to nine exons (*CcMADS23*), compared to the eight exons in *AtAP1*. *CcMADS70* (V2_13g015610.1) was annotated with seven exons, although we identified an eighth exon, leading to “*CcMADS70_corrected*”, coding a 211 a.a. protein with high sequence similarity to *LsAP1* ^[41]^ (Supplementary Figure S4B). With respect to protein domains, only *CcMADS12* contained significant SRF and K-domains, whereas the remainder had only one of the two domains. The euAPETALA1 (euAP1) motif ^[58][59]^ was identified in *CcMADS70_corrected*, supporting an *AP1* identity for this gene (Supplementary Figure S4B), which was hence named “*CcAP1*”. The euFUL and *AGL79* lineages could not be clearly distinguished from each other in the phylogenetic study. With respect to motifs, complete C-terminal euFUL motifs were only identified in CcMADS23 and CcMADS42 and absent in the remainder proteins. Taken together these results indicate that there are two *FUL-*like genes in globe artichoke that code the correct motif although without complete MADS and K-domains. *CcMADS42* was designated “*CcFULLike-A*” and *CcMADS23* “*CcFULLike-B*”.

After this analysis we decided to use real-time PCR (RTqPCR) to assess the usefulness of the *CcSOC1a*, *CcSOC1b*, and *CcFULLike-B*, MADS-box genes as markers for the process of transition to the reproductive phase in globe artichoke. We again used plants from early-bolting genotype c1, and late-bolting one c154, grown under vernalization and no-vernalization conditions. Shoot apices were sampled at three different time points, before floral transition (day 67 after transplanting), day 88 after transplanting (average time for floral transition in non-vernalized c1plants (Table 2)) and for c154 at a third time point at day 130 post transplanting (average time for floral transition in non-vernalized c154 plants (Table 2)) (Figure 5).

**Figure 5.**
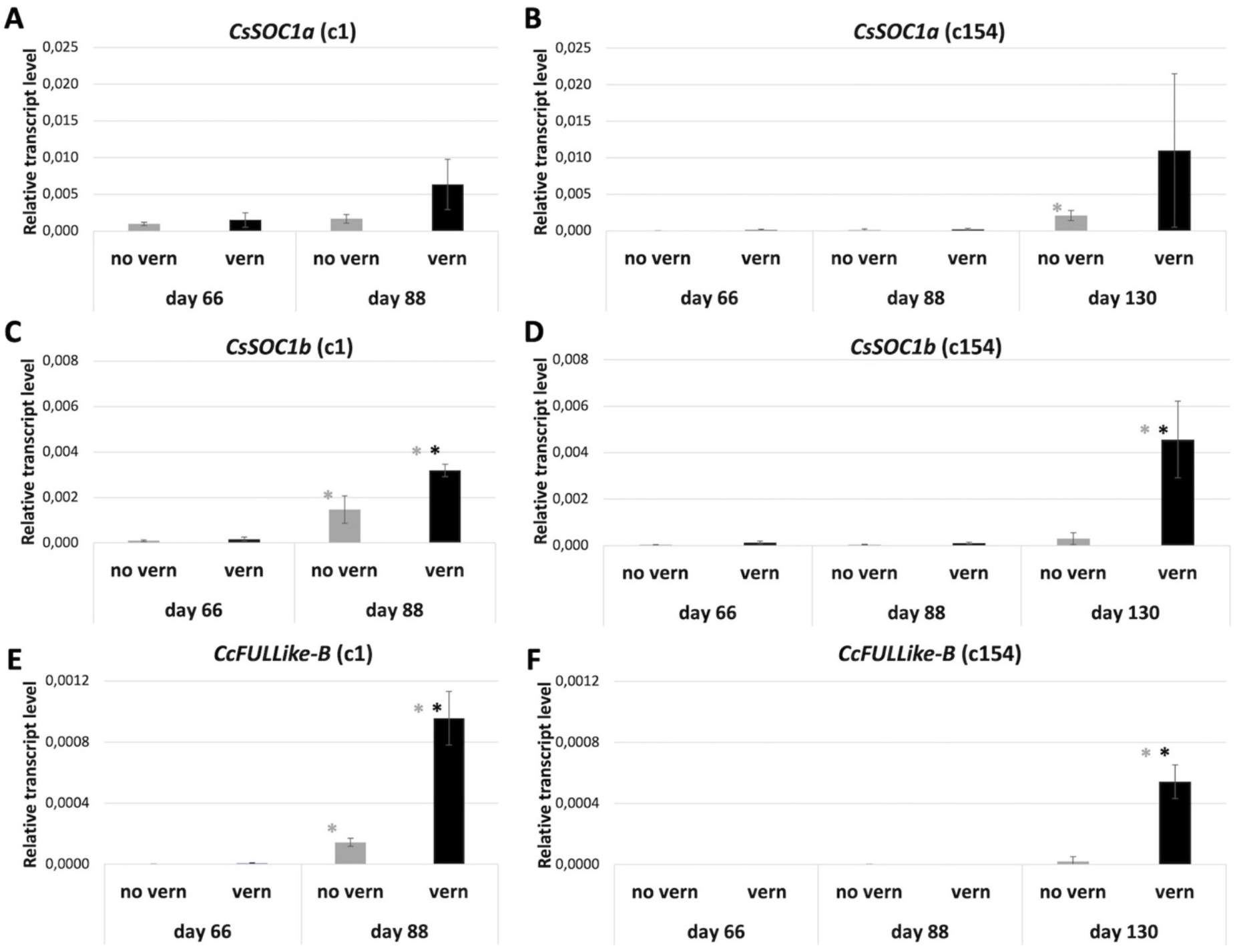
Expression of CcSOC1a, CcSOC1b and CcFULLike-B genes in early- and late-bolting genotypes. Expression level for *CcSOC1a* (A, B), *CcSOC1b* (C, D) and *CcFULLike-B* (E, F) genes was determined in samples taken before the floral transition and at pre-bolting stage 1. Analysis was performed on plants from the early-bolting genotype c1 (A, C, E) and the late-bolting genotype c154 (B, D, F) that have gone through vernalization or not. Error bars represent standard deviations of three samples. Black asterisks indicate significant differences (P ≤ 0.05) between treatments. Gray asterisks indicate significant differences (P ≤ 0.05) respect to the previous time-point. Genotype c1 was not sampled at day 130.

In agreement with the inferred roles for these genes, expression of *CcSOC1a*, *CcSOC1b*, and *CcFULLike-B* was very low or absent in apices in pre-bolting stage 0 before the floral transition (day 67), independent of growing conditions (Figure 5 A-F). At 88 days, when the floral transition had started in genotype c1 under both vernalization and no vernalization conditions, we detected a clear up-regulation of *CcSOC1b*, and *CcFULLike-B* genes respect to the values of day 67, being stronger in the plants that were in vernalization conditions (Figure 5 C, E), while the CcSOC1a expression did not change. On the other hand, at this same time point, no change in gene expression was detected in genotype c154 for none of the three tested genes, independent of growing conditions (Figure 5 B, D, F). At 130 days, expression of *CcSOC1b*, and *CcFULLike-B* genes was clearly detected in shoot apices of vernalized c154 plants, whereas expression did not change respect to the previous measurements of day 67 and 88 in the non-vernalized ones (Figure 5 D, F). Again, no clear changes in expression of *CcSOC1a* in the genotype c154 at day130 were observed. These results confirm that *CcSOC1b*, and *CcFULLike-B,* are strongly upregulated during the floral transition in globe artichoke, being even more evident in the late flowering genotype c154. This indicates that *CcSOC1b*, and *CcFULLike-B* up-regulation, and probably the floral transition, is accelerated by the vernalization treatment in both early and late-bolting genotypes. Finally, our results also support the use of *CcSOC1b*, and *CcFULLike-B* genes as expression markers for early transition to bolting in globe artichoke.

## Discussion

Earliness is a key trait for globe artichoke breeders. Despite multiple studies having addressed the timing of flowering in model plant species, reports on the genetic architecture of this trait in crop species remain relatively scarce. Amongst other factors, this can likely be attributed to the complexity of growing and handling plants of large size in controlled environmental conditions. Globe artichoke is a clear example of this. In this study we aimed to answer the question of whether bolting in globe artichoke is controlled by vernalization, and how cold exposure under natural growth conditions is important for different genotypes of artichoke selected by their early- or late-bolting phenotypes. To our knowledge this is the first time that an experiment has been performed in which globe artichoke plants have been observed under controlled vernalization temperatures during their whole life cycle. This enabled the study of bolting time between vernalization and no-vernalization treatments under comparable conditions. Moreover, our setting allows vernalization to follow a natural cycle, making the results and experimental design applicable for breeding programs as well.

In addition to observing bolting visually in the field, we characterized morphological changes in the apex that are associated with the vegetative-to-reproductive phase transition and linked this transition to the expression of homologs of key flowering regulators *SOC1* and *FUL*. The results of our vernalization experiments indicate that early-bolting genotypes have a significantly lower vernalization response than late-bolting genotypes, suggesting that a reduced requirement for vernalization is a relevant component of earliness in globe artichoke.

### Effect of vernalization on bolting

Regarding the effect of vernalization, we found that the 698-935 hours below 10°C advanced the time of bolting on average by 10.0 and 23.5 days for early- and late-bolting genotypes respectively. Although to our knowledge there exist no reports of other studies which allow for direct comparison of results, reports on the effect of either artificial or natural vernalization on bolting time provide some general trends. Artificial vernalization prior to transplanting, equivalating to about 500 chilling hours (hours < 10°C), reduced the time to bolting with 11 days in the late-bolting variety ’Orlando F1’ but had no significant effect on three other commercial hybrids ^[60]^. Therefore, in that study the effect of the artificial vernalization treatment on bolting time is lower than the 23.5 days we found for late-bolting genotypes. Another study on the effect of artificially vernalizing varieties prior to transplanting has been described by Rangarajan et al., (2000) ^[49]^, who reported a significant effect on early yield after treating plantlets for about 450 hours at 13°C in combination with late planting in spring, which meant that little to no natural vernalization was present. The effect was larger for the late-bolting variety ’Green Globe Improved’ than for early-bolting variety ’Imperial Staŕ. Another study by García and Cointry (2010) ^[61]^ failed to find an effect of vernalizing seeds, seedlings and plantlets of early-bolting variety ’Imperial Staŕ for 240 hours at 3°C, perhaps because of a lack of vernalization requirement in this variety.

The results from these studies are in line with our results insofar that the effect of vernalization is largely genotype-dependent. Although complete control of environmental conditions was out of the scope of this study, the experimental design that we developed addresses some of the challenges associated with field experiments or artificial vernalization, allowing for direct comparisons between genotypes of the effect of vernalization on bolting time. Moreover, the use of a heated net house with plastic insulation potentially offers an effective and affordable way for breeders of large-sized field crops to apply selection pressure against vernalization requirement in their programs.

### Vernalization requirement in globe artichoke

Vernalization has been reported to be a major factor that determines flowering time in globe artichoke ^[44]^ ^[48]^. Similarly, genotypes with an early-bolting phenotype could reflect the absence of a vernalization requirement ^[42][43]^. In species like Arabidopsis and rice, vernalization is a gradual and quantitative process that prepares the plant to respond to other cues (environmental or endogenous) that promote the floral transition ^[9]^ ^[14]^. In our experiments, cold depended on the prevailing weather in each season, which dictated not only the amount of accumulated chilling hours, but also when cold started to be perceived by the plants. Despite observing a clear vernalization response in terms of bolting in all the genotypes tested, probably we are not detecting a saturated vernalization response. Interestingly, when we monitor the floral transition by morphological changes in the SAM, we observed that early-bolting genotype c1 initiates the floral transition independently of vernalization whereas the same transition is significantly advanced in late-bolting genotype c154 under vernalization conditions. This observation suggests that early-bolting genotypes do not require vernalization to initiate the floral transition, but that somehow low temperatures are able to modulate bolting, as differences in bolting time were observed for this genotype. At the same time, our analysis supports the view that vernalizing temperatures can advance the floral transition in late-bolting genotypes, as well as modulate bolting progression. As commented, our microscopic observations indicate that early-bolting genotypes, or at least the c1 genotype, initiate the floral transition independently of chilling temperatures. Interestingly, at the molecular level, we clearly detected a stronger activation of homologs of floral integrators *CcSOC1b*, and *CcFULLike-B*, in the vernalized c1 plants than in non-vernalized ones at the floral transition/pre-bolting stage 1, suggesting an early floral transition or a stronger activation of those genes, and hence vernalization response.

### Effect of photoperiod on bolting

When DTB is compared between genotypes and seasons it is apparent that late-bolting genotypes c20, c154 and CARI, under vernalization conditions, bolt within a relatively narrow time frame of 161-172 days in each season. Contrarily, when observed in no-vernalization conditions, DTB is more variable in these genotypes with a range of 177-211 days (Table I). This suggests that vernalized plants react more uniformly to a stable bolting cue, such as photoperiod or age, whilst non-vernalized plants present a later and more variable response to those possible cues. Photoperiod measured for vernalized late-bolting genotypes was around 11.5-11.9 hours around the moment of reaching bolting stage B. This may support the hypothesis that late-bolting genotypes are long-day plants. However, early-bolting genotypes c1 and c70, independent of the vernalization treatment, presented mean DTB values that varied 13-15 days between seasons (Table 1) and with a critical photoperiod measured around 9.6 hours, rendering the early-bolting genotypes short-day plants. These observations are in line with reports that suggest critical photoperiod in globe artichoke being genotype-dependent ^[42][43]^ hence putting emphasis on the genotype dimension when dealing with this trait. Our morphological analyses of the apex around the moment of the floral transition suggested that, at least for the early- and late-bolting genotypes tested, the floral transition took place in short-day conditions. These observations render globe artichoke a short-day plant in terms of floral transition, despite bolting occurring in both long and short-day conditions.

### No indications for devernalization or epigenetic memory of the vernalized state

In globe artichoke there have been mentions of the concept of devernalization as a heat-induced reversal of the vernalized state. Critical temperatures for this to occur however appear to vary widely according to different reports, with mentions being >18.3°C (65°F) ^[52]^, >26°C ^[48]^, >32°C ^[49]^, and >33°C ^[60]^. If temperatures above 28°C are taken as a measure for devernalization, these conditions were met for 170-240 hours in both the vernalization and non-vernalization compartments during our experiments. Most of these hours were however accumulated in the first month of the experiment, when vernalizing temperatures still had to occur, and in the last month, when most plants had already bolted.

Four of the varieties in this study were clones and this might beg the question of whether an epigenetic memory of vernalization ^[62]^ could have existed to explain the earliness in early-bolting clonal varieties. According to our results this appears not to be the case. Both early- and late-bolting clones reacted to the no-vernalization treatment, rendering a vernalized state prior to the experiments unlikely. Likewise, plants that have been successfully vernalized in the first season do not bolt earlier after resprouting in the next season and require being vernalized again ^[3][63]^.

### Morphological and molecular markers for the vegetative-to-reproductive phase transition

Different scales that describe phenological or developmental stages in artichoke have been published. A macroscopic 8-stage scale of the apex that spans the vegetative-to-reproductive phase change till anthesis of peripheral flowers was described by Foury (1967) ^[5]^ ^[8]^. Another scale with 15 stages by Baggio et al. (2011) ^[7]^ is specifically tailored to the development of the head from the moment it reaches about 1.3 cm in size until the end of the reproductive phase. A scale describing phenological stages in *Cyncara cardunculus* according to the Biologische Bundesanstalt, Bundessortenamt und CHemische Industrie (BBCH) scale has been developed as well ^[6]^. None of the scales specifically conveys the changes in the vegetative or inflorescence apex that occur between the vegetative-to-reproductive phase transition and the start of bolting, when the primary inflorescence becomes visible in the rosette. We addressed this by the addition of five pre-bolting stages to the scale originally published by Foury (1967) ^[5]^. For lettuce, stages of development in the apex have been reported, e.g. by Chen et al. (2018) ^[64]^. Where, for lettuce the scale is largely based on the development of an inflorescence with multiple capitulae, in globe artichoke the transitions are defined by a more pronounced order in which first the primary and secondary capitulae develop.

The morphological markers developed in this study are useful to place the beginning of the vegetative-to-reproductive phase transition in globe artichoke, either just before, or during, pre-bolting stage 1. This is the stage where shoot apical meristem doming occurs, which is also an indicator of the vegetative-to-reproductive phase transition in other plant species ^[65]^. In addition, we observed that pre-bolting stage 1 coincides with an upregulation of expression of *CcSOC1b* and *CcFULLike-B,* globe artichoke homologs of *SOC1* and *FUL*. This further supports the placement of the vegetative-to-reproductive phase transition at the start of pre-bolting stage 1. The connection between doming and the vegetative-to-reproductive phase transition however may not be universal for Asteraceae and instead in lettuce this transition has been placed after doming and at the start of the elongation phase ^[64]^.

### MADS box genes in globe artichoke

Phylogenetic analysis allowed us to identify 82 putative MADS-box genes in the artichoke genome and determine as well putative orthologs of genes in this family that are key elements in the control of flowering time in other plant species. In that regard, our focus has been on *FUL* and *SOC1* orthologs. *FUL* is known to act redundantly with *SOC1* in promoting flowering and the expression levels of both genes increase at the time of vegetative-to-reproductive phase transition in Arabidopsis ^[66]^ ^[67]^. This is in line with the marked increase in expression levels observed for *CcSOC1b* and *CcFULLike-B* around the estimated time of floral transition in globe artichoke. *SOC1* homologs with increased expression around the time of vegetative-to-reproductive transition have also been reported for other Asteraceae species such as Chrysanthemum^[39]^ ^[68]^ and lettuce ^[41]^.

Research on the role of FUL genes in Asteraceae is limited. Ectopic expression of two homologs of *FUL* from *Chrysanthemum morifolium* in tobacco led to early flowering ^[69]^. Overexpression of *FUL* ortholog *GSQUA2* from gerbera led to accelerated flowering in the same species ^[70]^. These results suggest that the function of *FUL* orthologues as flowering promotors is conserved in Asteraceae.

*FLC* is a central gene in the control of vernalization in Arabidopsis and Brassicaceae ^[71][72]^. Our phylogenetic study revealed that the FLC subfamily in globe artichoke contains two members (Supplementary Table S4). Of the two, *CcMADS65* is most likely a closer homolog of Arabidopsis *FLM* and *FLC* genes than *CcMADS10*, although both have low protein similarity to *FLC* from Arabidopsis (30.3% and 25.7% respectively). Furthermore, there exist few reports of *FLC* orthologues in Asteraceae. A MADS box gene in chicory (*Cichorium intybus*), with high protein sequence similarity to *FLC* from Arabidopsis, is repressed by vernalization and acts as a flowering repressor ^[32]^. Puglia et al., 2016 ^[73]^ report on the isolation of the partial sequence of *ccMFL*, an *FLC* homolog from cultivated cardoon that was however not further characterized. It is possible that *FLC* genes have been subject to neofunctionalization in Asteraceae, as suggested by a recent study of MIKC^C^ genes in *Chrysanthemum lavandulifolium*, where *FLC* genes might rather be involved in flower or capitulum development ^[74]^. The absence of a clear homologue of Arabidopsis *FLC* in globe artichoke could mean that vernalization requirement and response in this species are underlain by a different genetic architecture.

Since earliness is a key breeding target for commercial hybrids, gaining an understanding of the genetic architecture of this trait supports the development of tools for efficient selection and introgression into the most market-relevant elite genetic backgrounds. With the advent of affordable mass sequencing and data analysis, a promising approach towards identification of genes involved in earliness, and vernalization requirement, is transcriptome sequencing analysis. Under this approach, molecular markers for the floral transition, such as the ones developed in this study, will be a helpful tool to link genetic markers to bolting time and earliness.

Limitations of the study One limitation of our study was that only minimum temperatures were controlled, but not mean or maximum temperatures. This is due to the impracticality of raising large plants such as globe artichoke in the fully controlled environment of a phytotron. Since the net house compartment in which the non-vernalization treatment was applied required the construction of plastic walls for the heating the be effective, control of day time temperature was limited to manually raising or lowering the internal and/or outward-facing compartment walls to manage air exchange. This made temperature control challenging under high irradiance, high outside temperature, and low wind conditions and might have induced stress in the non-vernalized plants that could not be accounted for. Another limitation is the destructive nature of sampling shoot apices, which precluded multiple observations of gene expression in individual plants.

## STAR Methods

## RESOURCE AVAILABILITY

Further information and requests for resources and reagents should be directed to and will be fulfilled by the lead contact, Vicente Balanzá Pérez

## Materials availability

This study did not generate new or unique reagents.

### Data and code availability

- All data reported in this paper will be shared by the lead contact upon request.
- This paper does not report original code.
- Any additional information required to reanalyze the data reported in this paper is available from the lead contact upon request.

## EXPERIMENTAL MODEL AND STUDY PARTICIPANT DETAILS

### Plant material

A total of seven genotypes were selected from the BASF Vegetable Seeds artichoke breeding program. These constituted two early-bolting clones, “c1” and “c70”, two late-bolting clones, “c20” and “c154”, two early-bolting lines “VER” and “VESB” and one late-bolting line “CARI”. Clones were produced at the BASF|Nunhems Cell Biology Services lab in Haelen, the Netherlands, according to internal standard protocol. Care was taken to avoid temperatures <13°C during production and transport. Clones were delivered at the 3-4 leaf stage between 20 August and 31 August in the years 2019, 2020 and 2021. Lines were sown in the third week of July in a commercial nursery and allowed to germinate for seven days at 16°C. Both clones and lines were kept in pots under shade cloth for 1-2 weeks between delivery and the start of the experiments at the 4-5 leaf stage.

## METHOD DETAILS

### Experimental design and treatment

The experiments took place in a net house in vicinity of Águilas, Spain (230 m MSL). The limestone soil was solarized, tilled, and fertilized, with compost prior to transplanting, which took place on 25 September 2019, 25 September 2020 and 01 October 2021. Plants were assigned to plots of 10 individuals each, which were subsequently randomized, with experimental groups containing 30-70 individuals in accordance with availability. Planting distance was 0.8 m within the row and 1.5 meter between the rows. A border row made up of commercial hybrids surrounded the perimeter (Supplementary Figure S1).

Along the external wall of the net house, two compartments were created. In one compartment minimum temperatures were controlled by a thermostat-controlled oil heater set to 13°C. To allow the system to heat effectively, a double ceiling, an external wall, and internal walls were constructed from thin transparent plastic. The internal and external walls were raised in the morning by the grower and lowered in the late afternoon or early evening to allow for ventilation and to moderate daytime temperatures. The other compartment was identical, but with no heating system, double ceiling nor outward facing plastic wall, allowing circulation of cool air during the night. The temperature in each compartment was monitored with a HOBO MX2302A data logger with RS-3B solar radiation shields (both from Onset Computer Corp, Bourne, USA) mounted in the center of the compartment at a height of 1.5 m. Daylength was calculated with the R-package chillR ^[75]^ for a latitude of 37.49°N.

### Macroscopic characterization

Plants were observed weekly for symptoms of bolting according to the developmental scale we designed (Supplementary Table S1). Plants with multiple rosettes or suspected disease symptoms were removed from the experiment. Bolting observation campaigns lasted from 2019-05-10 until 2020-05-08 (150 days) during the 2019-2020 season, from 2020-11-30 till 2021-05-19 (170 days) and from 2021-12-13 till 2022-04-20 (128 days). The end dates for the bolting observation campaigns coincided with rising temperatures in the net house becoming unconducive to plant growth and development. Plants were considered to have bolted upon reaching bolting stage B and the number of days-to-bolting (DTB) was calculated for statistical analysis.

### Tissue fixation, sectioning and staining

At predetermined intervals, individuals randomly selected from genotypes were dissected and their apices were scored according to the developmental scale (Figure 1, Supplementary Table S1). During the 2021-2022 season, apices were preserved for microscopy by fixation for 24h in a buffer consisting of 50% (v/v) ethanol, 10% formaldehyde and 5% (v/v) acetic acid, followed by dehydration in ethanol and storage at 4°C. For microscopy, the samples were infiltrated and embedded in paraffin and cut with a microtome to 8 μm sections. Toluidine staining was performed according to the protocol described by ^[76]^. Microscopy was performed using a Leica 5000 microscope (Leica, Germany).

### Identification of MADS box genes in globe artichoke

Protein sequences from MADS box genes were obtained for Arabidopsis (The Arabidopsis Information Resource ^[77]^) ^[78]^, tomato (Sol Genomics Network ^[79]^), safflower ^[37]^, *Chrysanthemum nankingense* ^[80]^, lettuce^[35]^ and gerbera ^[70]^. For lettuce, an additional AP1 homologue was included from ^[41]^. Arabidopsis MADS box genes were BLASTed against the 28.632 predicted proteins in globe artichoke ^[55]^. In the resulting 82 hits, Pfam SRF (PF00319.21) and K-box (PF01486.20) domains were predicted by a Hidden Markov model using HMMER 3.1b1. Protein sequences were aligned with MUSCLE ^[81]^. Maximum likelihood (ML) phylogenies were inferred with the Phangorn v2.10 package. The optimal model was selected modelTest, followed by 1000 iterations of Nearest Neighbor Interchange (NNI) bootstraps. Phylograms were constructed and annotated with ggtree ^[82]^.

### RNA sampling and Q-PCR analysis

SAMs dissected in the net house were immediately transferred to 300 μl RNAlater™ solution (Thermo Fisher Scientific, USA) for storage at -18°C. RNA extraction was performed with the EZNA® Plant RNA kit (Omega Bio-tek, USA) with on-column DNAse treatment with an RNase-free DNase I Set (Omega BioTek, USA) and converted to cDNA with a SuperScript IV First-Strand Synthesis System (Thermo Fisher, USA). Primers were designed for *CcSOC1a*, *CcSOC1b*, *CcAP1* and *CcFULLike-A* and *CcFULLike-B* with CLC Genomics Workbench v21.0.3 (QIAGEN, Aarhus, Denmark) (Supplementary Table S5). Amplicons were checked with high resolution melting prior to qPCR on a magnetic induction cycler (Mic) qPCR system (Bio Molecular Systems, Australia). Relative expression was calculated according to the 2^!Δ#$^ method. From *CcAP1* and *CcFULLike-A* no pure amplicon could be obtained precluding quantification of these two genes.

## QUANTIFICATION AND STATISTICAL ANALYSIS

### Statistical analysis

Bolting data were modelled using a linear mixed model with the main effects (genotype, treatment and season) and the interaction terms (genotype: treatment and treatment:season) set as fixed effects. The term “plot” was set as a random factor and nested within “season” to account for the between plot variation within each season. The estimated means were extracted from the model for each “genotype:treatment” combination, and the significance of differences between best linear unbiased estimators (BLUEs) was assessed by calculating the LSD and determining whether the difference between BLUEs was larger or smaller than the value of the LSD (6.86 days). In an additional modelling to determine the effect of the treatment on early-versus late-bolting genotypes, the explanatory variable “genotype” was substituted with “earliness”, a dichotomous variable distinguishing early (genotypes c1, c70, VESB and VER) from late (genotypes c20, c154 and CARI). DTpb1values for genotypes c1 and c154 were analyzed with a one-sided Student’s t-test and qPCR results with a two-sided t-test, both with α = 0.05.

Floral transition data were analyzed with a two samples t-test, one tail, considering p-value<0.05 as significant.

## Acknowledgements

The authors like to thank BASF | Nunhems for the support and facilities that made the experiments possible. In particular, the artichoke breeding team, the Statistics team, and external contactors, have been crucial to this work. Michael Dobres (Head of Breeding EMEA II at BASF | Nunhems at the time) and Peter Visser (R&D Crop Lead for leafies, artichoke, okra, celeriac at BASF | Nunhems) facilitated the collaboration project between BASF | Nunhems and CSIC-IBMCP.

## Authors’ contributions

- Conceptualization and methodology: P.V., F.M., V.B., Re.B. and Ri.B.
- Investigation: Ri.B, Re.B., and V.B.
- Writing – Original Draft: Ri.B.
- Writing – Review & Editing: Ri.B., V.B. and F.M.
- Funding Acquisition: F.M. and P.V.
- Supervision, P.M., V.B. and Re.B.

## Supplementary Figures

**Supplementary Figure S1:**
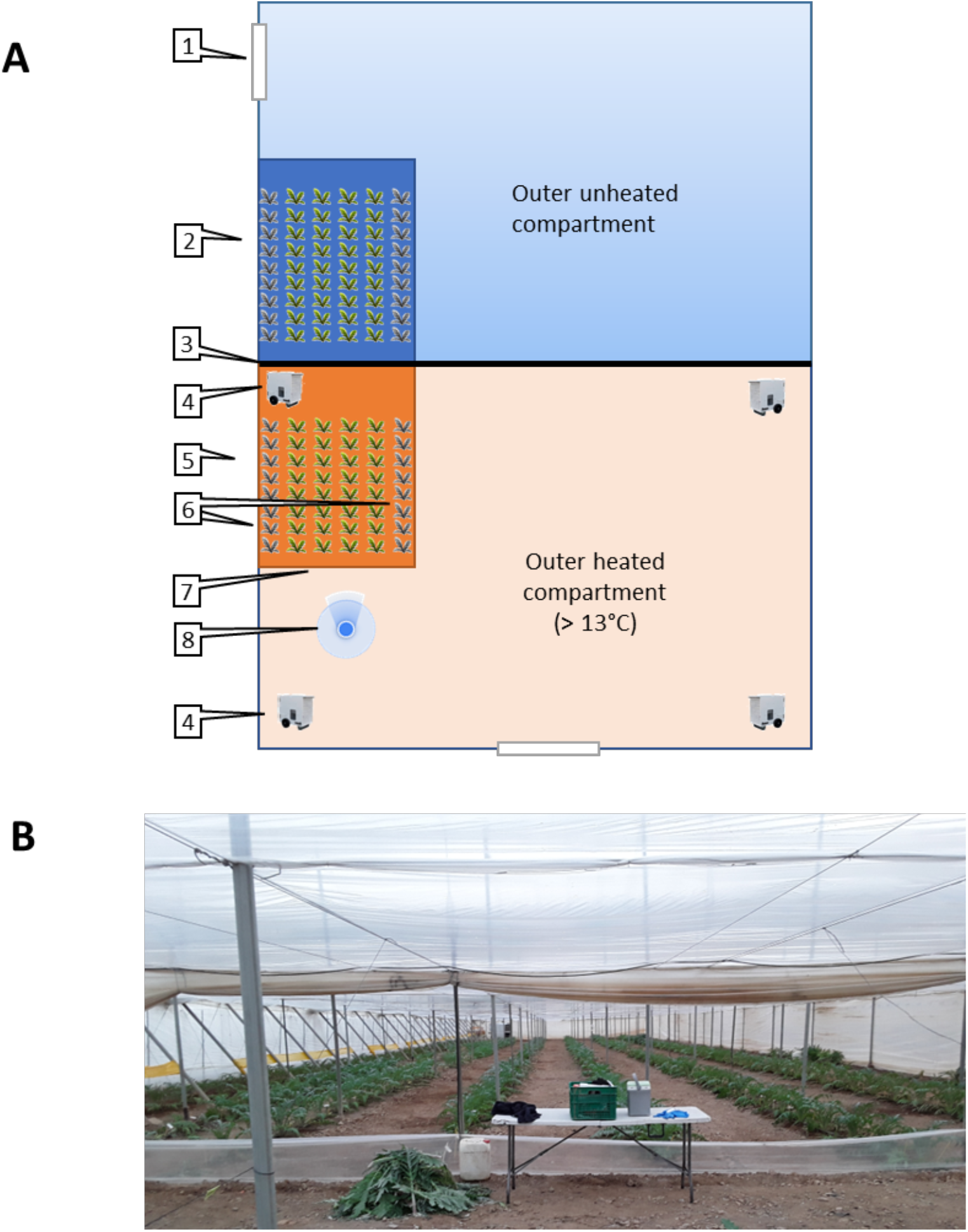
Schematic representation and photographic illustration of experimental conditions. **A.** Schematic representation of the experiment. 1 = access door in outer wall, 2 = cold compartment (vernalization / control) (blue fill), 3 = double wall separating heated and non-heated sectors, 4 = heater, 5 = heated compartment (red fill), 6 = border rows (grey), 7 = inner compartment wall, 8 = viewpoint for the photograph below. **B.** Photograph of the heated compartment from the viewpoint indicated by marker (8). The inner compartment wall facing the camera has been raised while the lateral inner wall and outer wall are still lowered for comparison. In the back of the image is the double wall separating the sectors, behind which is the cold compartment. Yellow bands along the outer wall serve as pest control. The table and tools illustrate the scale of the image.

**Supplementary Figure S2:**
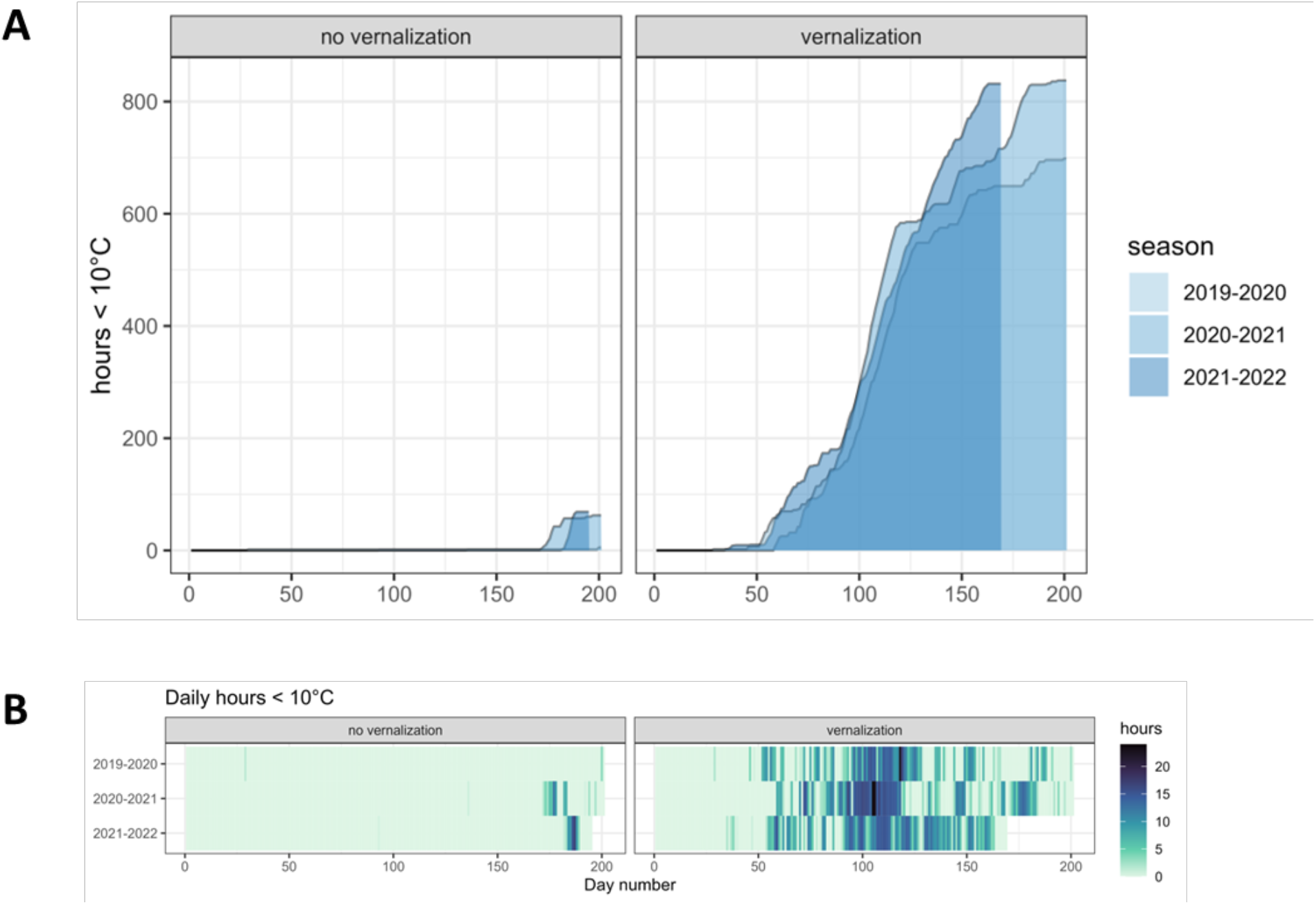
Accumulation and distribution of hours < 10°C. **A** Accumulated numbers of hours <10°C for each season in the no-vernalization and vernalization compartments. **B** Number of hours <10°C for each individual day. Data from three seasons.

**Supplementary Figure S3:**
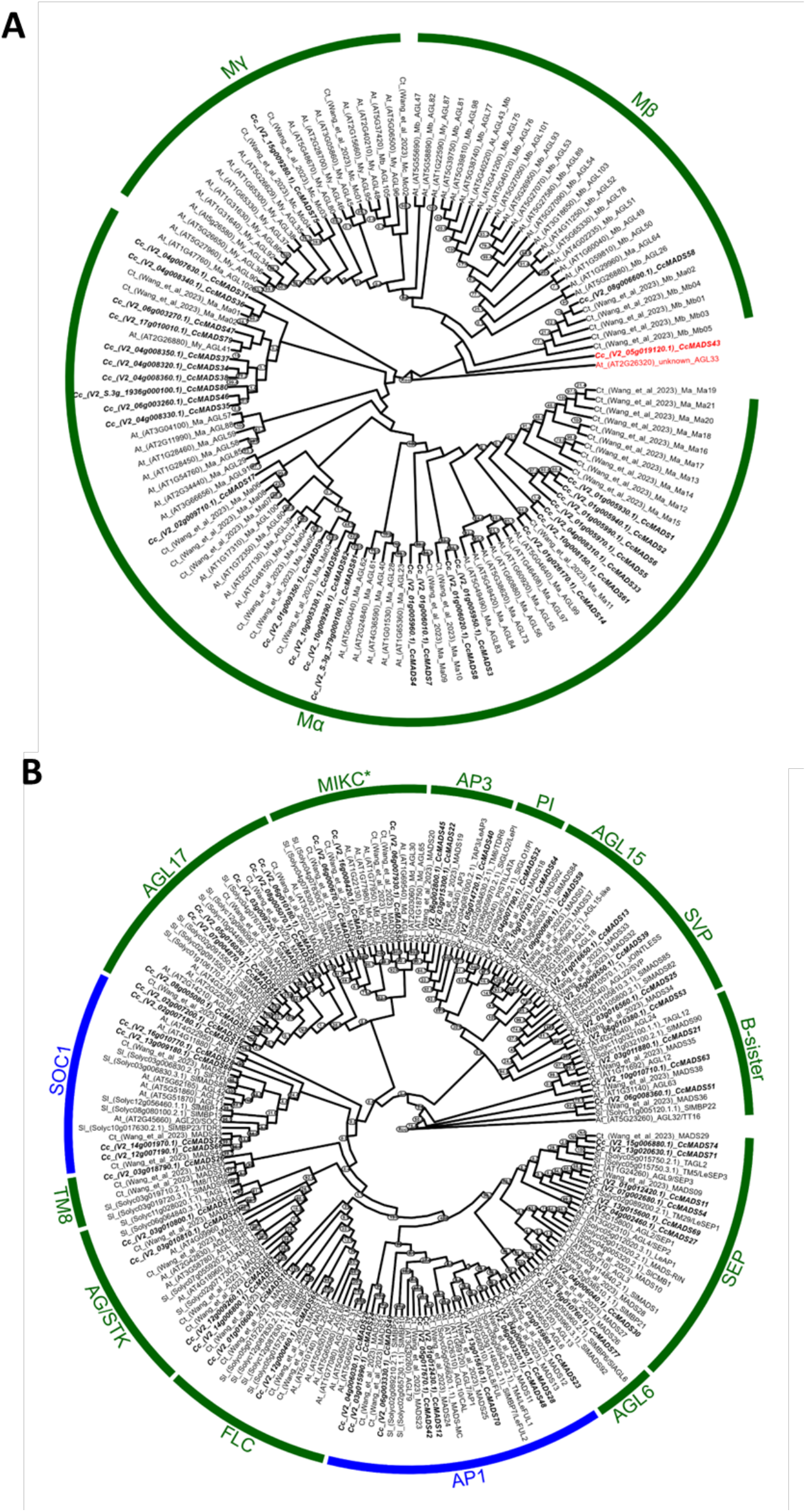
ML cladograms of Type I and Type II MADS box genes. **A** Type I MADS box genes from globe artichoke and Arabidopsis. Red tip names indicate ungrouped genes. **B** Type II MADS box genes from globe artichoke (Cc), Arabidopsis (At), tomato (Sl) and safflower (Ct). Green strips annotate subclades for both Type I and Type II cladograms, with the AP1 and SOC1 subclades highlighted in blue.

**Supplementary Figure S4:**
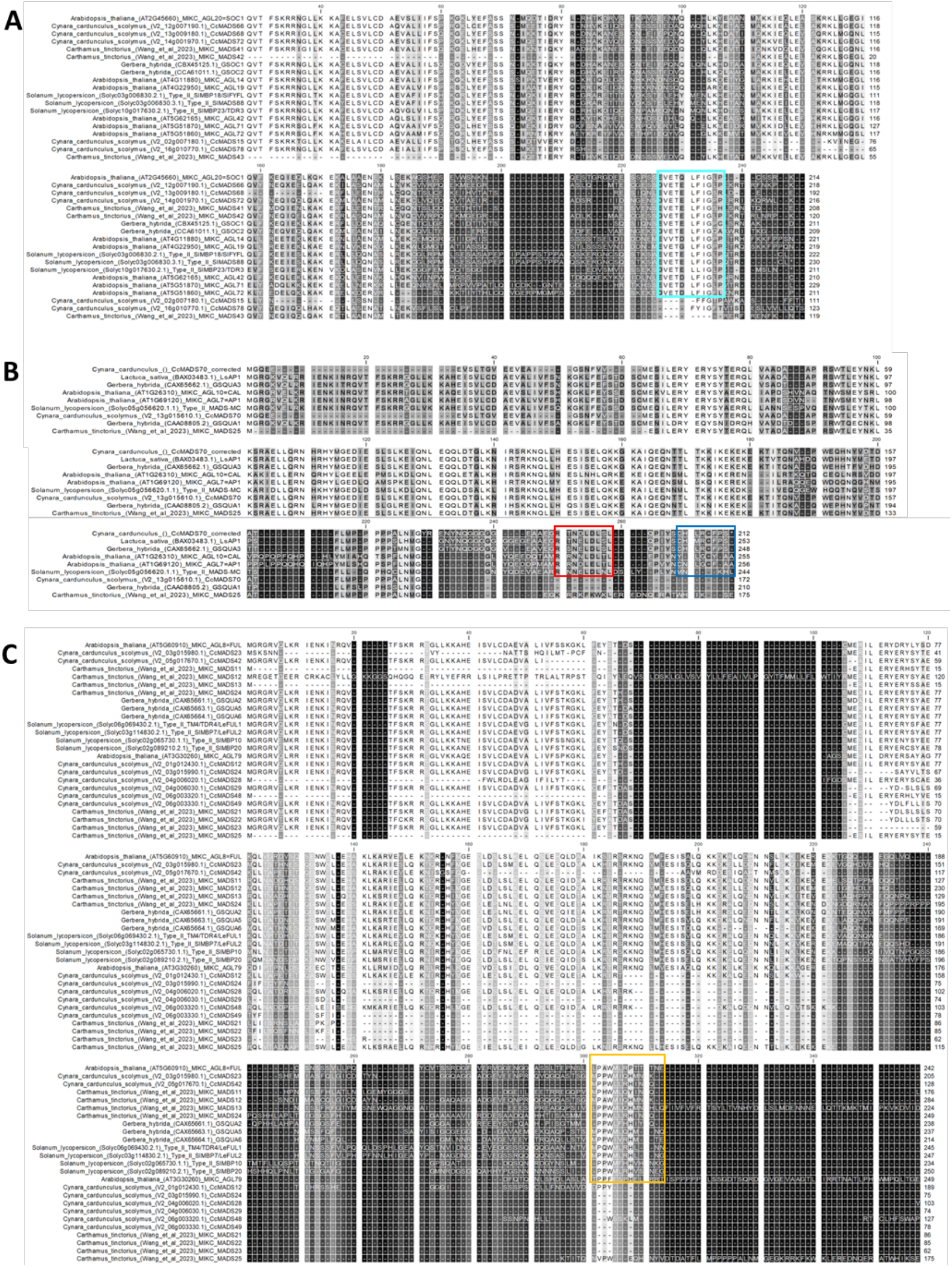
Protein sequence alignments for SOC1, AP1 and FUL subfamilies. Genes used in the phylogenetic study. **A** SOC1 subfamily, TM3/SOC1 motifs marked with a cyan box. **B** AP1 subfamily. Red box denotes the euAP1 acidic domain and the blue box the farnesylation motif. C FUL subfamily. The euFUL/paleoAP1 domain is marked by a yellow box.

## Supplementary tables

**Supplementary Table S1:**
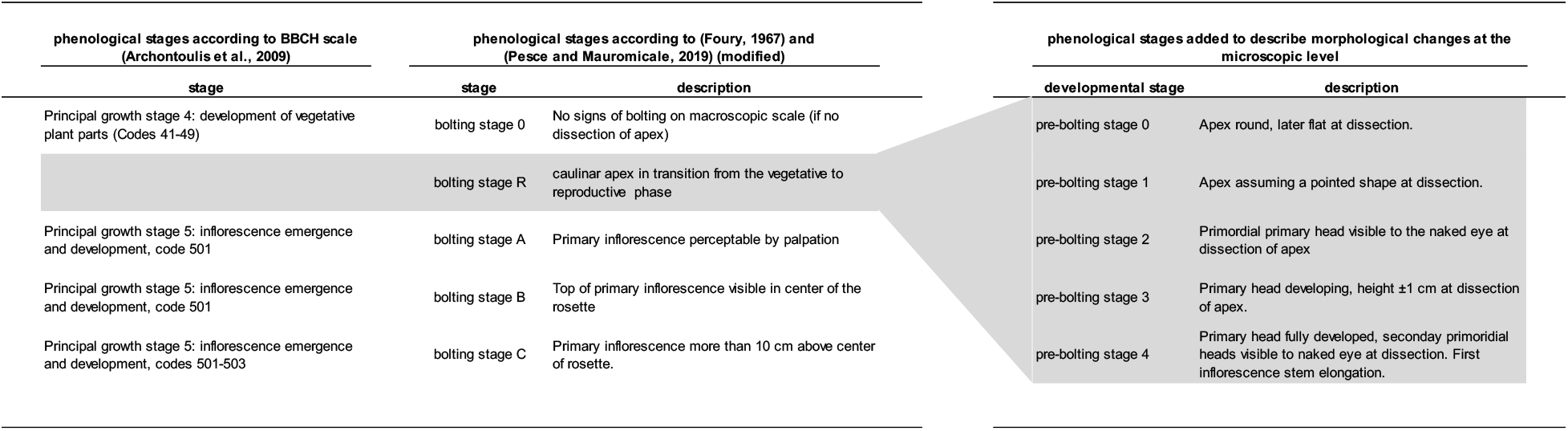
Comparison between phenological stages around the time of floral transition and the stages added to describe morphological changes. Grey fill accentuates the positioning of the added bolting stages in relation to the existing scales.

**Supplementary Table S2:** Environmental data calculated and collected during the experiments.

| season | treatment | n_days | daily mean (°C) |  |  | accumulated |
| --- | --- | --- | --- | --- | --- | --- |
|  |  |  | min | mean | max | hours<br>< 10°C |
| 2019-2020 | no vernalization | 201 | 19.1 | 14.4 | 28.3 | 5 |
| 2020-2021 | no vernalization | 201 | 18.1 | 13.4 | 27.4 | 62 |
| 2021-2022 | no vernalization | 195 | 17.6 | 13.6 | 26.2 | 69 |
| 2019-2020 | vernalization | 201 | 16.7 | 10.6 | 27.5 | 698 |
| 2020-2021 | vernalization | 201 | 16.1 | 9.9 | 26.8 | 837 |
| 2021-2022 | vernalization | 169 | 15.8 | 9.9 | 27.5 | 832 |

**Supplementary Table S3:** Wald tests for non-zero predictors. Wald tests for models per genotype (genotypes c1, c70, c20, and c154) and per category (early-bolting genotypes c1 and c70 vs late-bolting genotypes c20 and c154).

| model<sup>a</sup> | source | Df<sup>b</sup> | denDF<sup>c</sup> | F.inc<sup>d</sup> | Pr<sup>e</sup> |
| --- | --- | --- | --- | --- | --- |
| **per genotype** | (Intercept) | 1 | 100.6 | 94810 | 1.85E-151 |
|  | genotype | 6 | 107.3 | 615.1 | 1.16E-80 |
|  | treatment | 1 | 99.8 | 271.8 | 3.01E-30 |
|  | season | 2 | 51.6 | 1.537 | 2.25E-01 |
|  | genotype:treatment | 6 | 106.8 | 6.383 | 9.09E-06 |
|  | season:treatment | 2 | 52.5 | 7.253 | 1.66E-03 |
| **per category** | (Intercept) | 1 | 154.7 | 24030 | 1.35E-171 |
|  | earliness | 1 | 147.7 | 718.6 | 1.29E-58 |
|  | treatment | 1 | 154.6 | 48.21 | 9.95E-11 |
|  | season | 2 | 74.2 | 0.3074 | 7.36E-01 |
|  | earliness:treatment | 1 | 146.9 | 14.47 | 2.08E-04 |
|  | treatment:season | 2 | 74.1 | 6.669 | 2.17E-03 |

<sup>a</sup> Model used: "per genotype", each genotype, "per category", early-bolting vs late-bolting genotypes
<sup>b</sup> Df: Degrees of freedom
<sup>c</sup> denDF: denominator degrees of freedom
<sup>d</sup> F.inc: F-statistic value
<sup>e</sup> Pr = p-value

**Supplementary Table S4:**
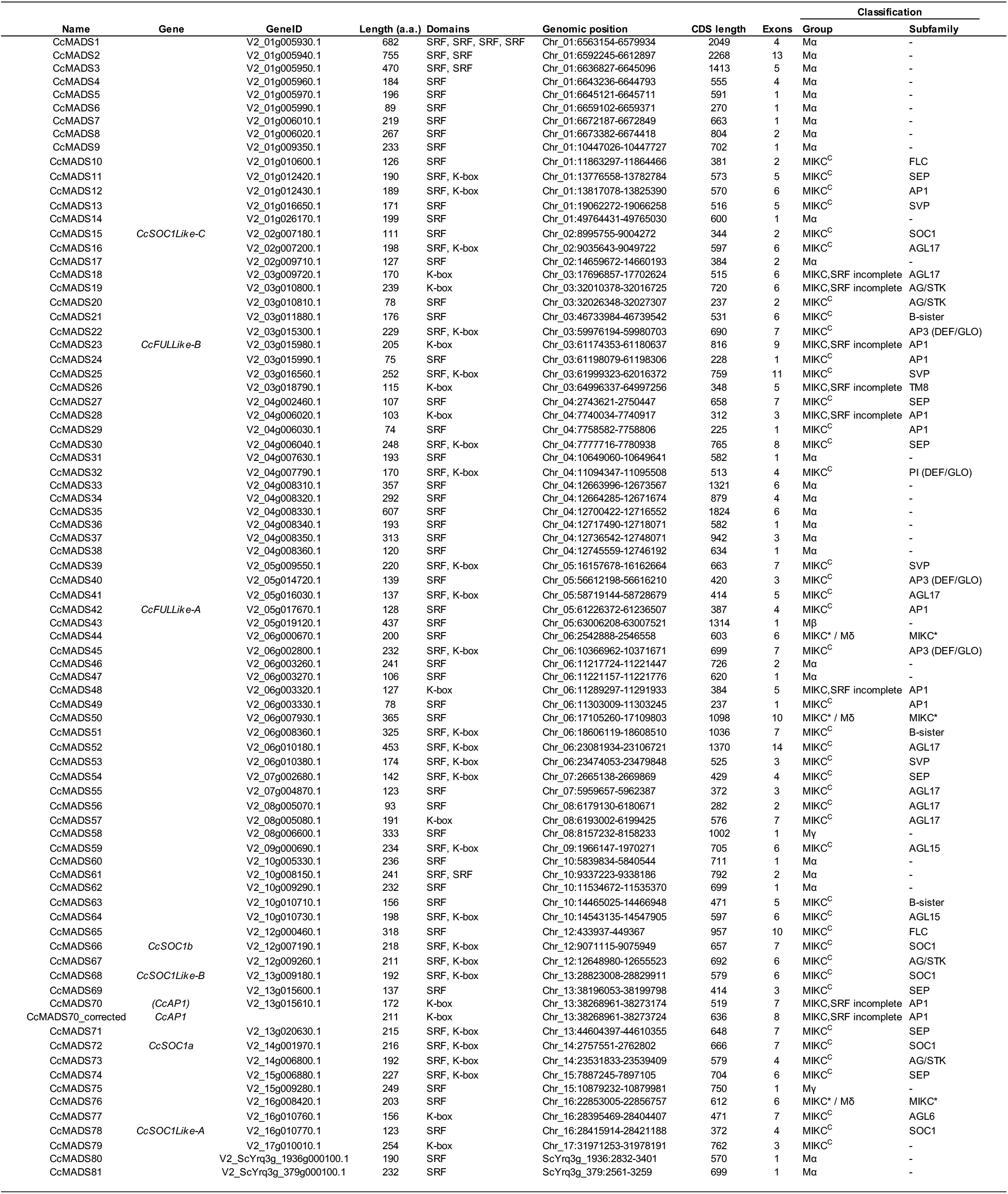
MADS box genes from globe artichoke.

**Supplementary Table S5:**
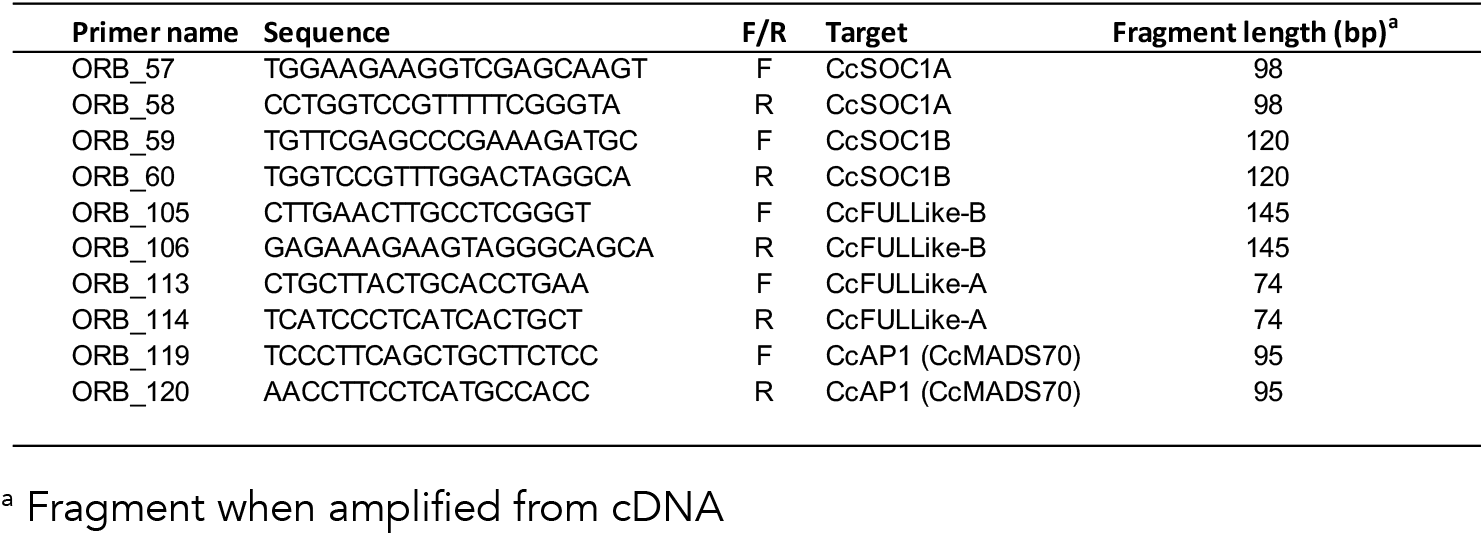
Primers developed.

